# Restoration of Redox Homeostasis and Endogenous Aldehyde Detoxification by UT-018 Following Acute Ethanol Exposure

**DOI:** 10.64898/2026.07.28.741198

**Authors:** Uday Saxena, Sadiya Mehboob, Shreya Shahapur, Tanisha Samal, Priyanka Jhadav, Gopi Kadiyala, Markandeya Gorantla

## Abstract

Alcohol-induced toxicity is driven largely by the accumulation of acetaldehyde and disruption of hepatic redox homeostasis during ethanol metabolism. Oxidation of ethanol by alcohol dehydrogenase (ADH) consumes nicotinamide adenine dinucleotide (NAD) while generating NADH, shifting the intracellular redox state toward a highly reduced environment that impairs mitochondrial function, limits endogenous aldehyde dehydrogenase (ALDH)-mediated acetaldehyde clearance, and promotes oxidative stress and tissue injury. We investigated whether UT-018, a novel metabolic intervention, could support endogenous metabolic resilience during acute ethanol exposure using complementary in vitro and in vivo models. Mechanistic in vitro studies evaluated ADH-dependent NADH generation and NAD add-back experiments, while in vivo investigations assessed serum ALDH-associated activity, circulating acetaldehyde concentrations, and gross gastrointestinal and hepatic morphology following acute ethanol challenge. UT-018 reduced ethanol-associated NADH accumulation in a concentration-dependent manner without evidence of irreversible ADH inhibition. Restoration of NADH generation following supplementation with exogenous NAD demonstrated reversible modulation of ethanol-associated redox biology rather than direct enzymatic inhibition. In vivo, UT-018 enhanced serum ALDH-associated activity, reduced circulating acetaldehyde concentrations by approximately 27 to 33% compared with ethanol-treated controls. Metabolic biomarkers were accompanied by preservation of gross colon and liver morphology following acute ethanol exposure. Collectively, these findings support coordinated biological activity across multiple interconnected stages of alcohol metabolism and support a systems-level mechanism in which restoration of redox homeostasis enhances endogenous aldehyde detoxification, reduces acetaldehyde burden, and preserves tissue integrity. These results identify alcohol metabolism restoration as a promising strategy for enhancing physiological resilience to acute alcohol exposure and provide a rationale for further preclinical and clinical evaluation of UT-018.

**Highlights:**

- UT-018 restored ethanol-associated redox homeostasis by reducing excessive NADH accumulation without irreversible inhibition of alcohol dehydrogenase in vitro.
- Restoration of redox balance was associated with enhanced endogenous aldehyde dehydrogenase (ALDH)-associated activity following acute ethanol exposure in vivo.
- UT-018 reduced circulating acetaldehyde concentrations by approximately 30%.
- The metabolic homeostasis was accompanied by preservation of gross gastrointestinal and hepatic morphology in an acute ethanol challenge model.
- The collective findings support a systems-level mechanism in which modulation of endogenous alcohol related metabolic pathways enhances physiological resilience to acute alcohol exposure.

## Introduction

Alcohol consumption remains one of the leading preventable causes of liver disease, gastrointestinal injury, metabolic dysfunction, and premature mortality worldwide. Although ethanol itself exerts biological effects, increasing evidence indicates that acetaldehyde—a metabolic intermediate of ethanol oxidation is responsible for much of the cellular toxicity associated with alcohol exposure. Acetaldehyde readily forms adducts with proteins, nucleic acids, and membrane lipids, initiating oxidative stress, mitochondrial dysfunction, inflammation, and tissue injury.

The metabolism of ethanol is tightly linked to cellular redox biology. During oxidation of ethanol by alcohol dehydrogenase (ADH), NAD^+^ is reduced to NADH, increasing the intracellular NADH/NAD^+^ ratio. Excessive NADH accumulation perturbs β-oxidation, gluconeogenesis, mitochondrial respiration, and tricarboxylic acid (TCA) cycle activity, thereby reducing the efficiency of endogenous aldehyde detoxification.

Most currently available interventions intended to reduce alcohol-associated symptoms focus on hydration, hantioxidants, herbal extracts, or vitamins. While these approaches may alleviate symptoms, they generally do not address the metabolic disturbances responsible for acetaldehyde accumulation and redox imbalance. No currently available intervention directly targets restoration of alcohol-induced hepatic redox imbalance.

We hypothesized that coordinated support of endogenous metabolic pathways could provide a more physiologically relevant approach to alcohol detoxification. UT-018, a novel investigational metabolic intervention, was therefore evaluated for its effects on ethanol-associated redox modulation, endogenous aldehyde detoxification, serum acetaldehyde accumulation and tissue protection.

## Materials and Methods

### Study Design

The study consisted of complementary experimental components. Mechanistic in vitro studies evaluated the effect of UT-018 on alcohol dehydrogenase (ADH)-dependent NADH generation together with NAD^+^ add-back experiments to distinguish reversible modulation of redox homeostasis from irreversible enzyme inhibition. These findings were subsequently evaluated in vivo using an acute ethanol challenge model in which serum aldehyde dehydrogenase (ALDH)-associated activity, circulating acetaldehyde concentrations, and gross tissue morphology were assessed. The integrated experimental design enabled evaluation of sequential biological events extending from ethanol metabolism to tissue injury.

Primary study endpoints included:

- ADH-dependent NADH generation
- NAD^+^ add-back mechanistic studies
- Endogenous serum ALDH-associated activity
- Circulating acetaldehyde concentration
- Gross colon morphology
- Gross liver morphology

### Investigational Formulation

UT-018 is an investigational metabolic intervention developed to support endogenous physiological pathways involved in alcohol metabolism rather than functioning as an exogenous alcohol-metabolizing system.

### Animal Model

The in vivo investigation was performed in **male Sprague Dawley rats** (6–8 weeks of age) weighing approximately **200 ± 20 g** at study initiation. Animals were obtained from a CPCSEA-registered commercial breeder and were acclimatized for a minimum of seven days before commencement of dosing.

Animals were housed under controlled environmental conditions (temperature 19–25°C; relative humidity 30–70%; 12-hour light/dark cycle) with ad libitum access to standard laboratory chow and drinking water except during the scheduled fasting period preceding ethanol administration. Animals were fasted overnight (approximately 16 h) before dosing, food was returned four hours after dosing, and water was restored two hours after dosing. Each group consisted of eight animals. Test formulations were administered orally at a constant dosing volume of **10 mL/kg**.

### Ethical Approval

All animal experiments were performed in accordance with the guidelines of the Committee for Control and Supervision of Experiments on Animals (CCSEA), Government of India. Experimental procedures were apsupportd by the Institutional Animal Ethics Committee. The study was conducted at Cology Biosciences Pvt. Ltd., Hyderabad, India, under institutional animal care guidelines.

## Dose Preparation and Administration

A 100 mM UT-018 stock formulation was prepared. Lower concentrations were generated by dilution with Milli-Q water to obtain final concentrations of 25 mM and 50 mM, while the 100 mM formulation was administered directly. Comparator formulations were prepared according to sponsor specifications.

All investigational formulations were administered by oral gavage at a dose volume of **10 mL/kg**, calculated from the most recent individual body weight.

Ethanol was prepared as a 38% (v/v) solution from absolute ethanol and administered orally at 3 g/kg body weight one hour after administration of the assigned formulation. This dosing paradigm was selected to evaluate whether pretreatment with UT-018 altered the metabolic response to subsequent acute ethanol exposure.

## In Vitro Redox Assay

A recombinant alcohol dehydrogenase commercially available system was used to investigate the influence of UT- 018 on ethanol-associated redox metabolism. Reaction mixtures containing alcohol dehydrogenase, ethanol, NAD^+^, and increasing concentrations of UT-018 were incubated under standardized conditions. Formation of NADH was monitored spectrophotometrically as an indicator of ethanol oxidation.

To distinguish reversible metabolic modulation from irreversible enzyme inhibition, parallel reaction mixtures received supplemental NAD^+^ following incubation. Restoration of NADH generation after cofactor supplementation was interpreted as evidence that UT-018 preserved enzyme function while modulating the redox environment.

## Acute Ethanol Challenge

Following a one-week acclimatization period, animals received their assigned oral treatment. One hour later, ethanol was administered to all designated treatment groups.

Blood samples were collected before dosing and at **0.5, 1, 2, 4, and 6 hours** after ethanol administration. Approximately 0.25–0.40 mL of blood was collected at each time point by retro-orbital sampling. Samples were processed immediately for preparation of plasma and serum by centrifugation at **4**,**000 rpm for 10 minutes at 4°C**. Aliquots were stored at **−80°**C until biochemical analysis.

At six hours after ethanol administration, animals were euthanized by carbon dioxide inhalation. Colon and liver tissues were rapidly excised, photographed, snap-frozen and stored at **−80°C**. Serum and plasma samples were retained for subsequent biochemical analyses, including ALDH-associated activity and acetaldehyde measurements.

## Determination of Serum ALDH Activity

Serum samples collected following ethanol challenge were analyzed for endogenous ALDH-associated activity using validated biochemical methods. Measurements were obtained at serial time points to characterize the temporal relationship between UT-018 administration and endogenous aldehyde detoxification. The principal endpoint was enhancement of physiological ALDH activity relative to ethanol-treated controls.

## Determination of Circulating Acetaldehyde

Circulating acetaldehyde concentrations were quantified following acute ethanol administration using validated analytical methodology. Comparisons were made between vehicle-treated animals, UT-018-treated groups, and comparator groups to determine whether UT-018 enhanced endogenous acetaldehyde clearance.

## Gross Evaluation of Colon and Liver

Following euthanasia, the colon and liver were examined macroscopically for evidence of ethanol-associated injury. Gross observations included assessment of tissue color, congestion, edema, and overall structural integrity. Representative images were recorded for comparison among treatment groups.

Macroscopic findings were interpreted together with biochemical endpoints to evaluate whether modulation of ethanol metabolism translated into preservation of tissue morphology within the gut–liver axis.

## Clinical Observations

Animals were monitored throughout the study for mortality, morbidity, and treatment-related clinical signs. Cage-side observations were recorded at predetermined intervals throughout the six-hour observation period. No unexpected adverse effects attributable to UT-018 administration were reported during the study, and clinical observations were consistent with the expected transient effects of acute ethanol exposure.

## Statistical Analysis

Data are expressed as **mean ± standard error of the mean (SEM)** unless otherwise indicated. Statistical comparisons among treatment groups were performed using one-way or two-way analysis of variance (ANOVA), followed by an appropriate post hoc multiple-comparison procedure (e.g., Dunnett’s or Tukey’s test). Time-course analyses were evaluated using repeated-measures ANOVA where appropriate. A two-sided P value of <0.05 was considered statistically significant.

## Results

### UT-018 Modulates Ethanol-Associated Redox Biology in vitro

UT-018 reduced NADH accumulation in a concentration-dependent manner during ADH-mediated ethanol oxidation in vitro. Importantly, supplementation with exogenous NAD^+^ restored NADH production toward control values, indicating that UT-018 did not irreversibly inhibit alcohol dehydrogenase but instead modulated the ethanol-associated redox environment. Following incubation of recombinant ADH with ethanol and UT-018 (25 mM), excess NAD^+^ was added to parallel reaction mixtures. Restoration of NADH generation after supplementation with exogenous NAD^+^ indicates that UT-018 does not irreversibly inhibit alcohol dehydrogenase but instead reversibly modulates the intracellular redox environment. These findings support preservation of physiological ethanol metabolism while limiting excessive redox imbalance. NAD^+^ add-back experiments demonstrated reversible modulation of ethanol metabolism. This observation suggests that UT-018 supports support of intracellular redox homeostasis during ethanol metabolism.

**Figure 1.**
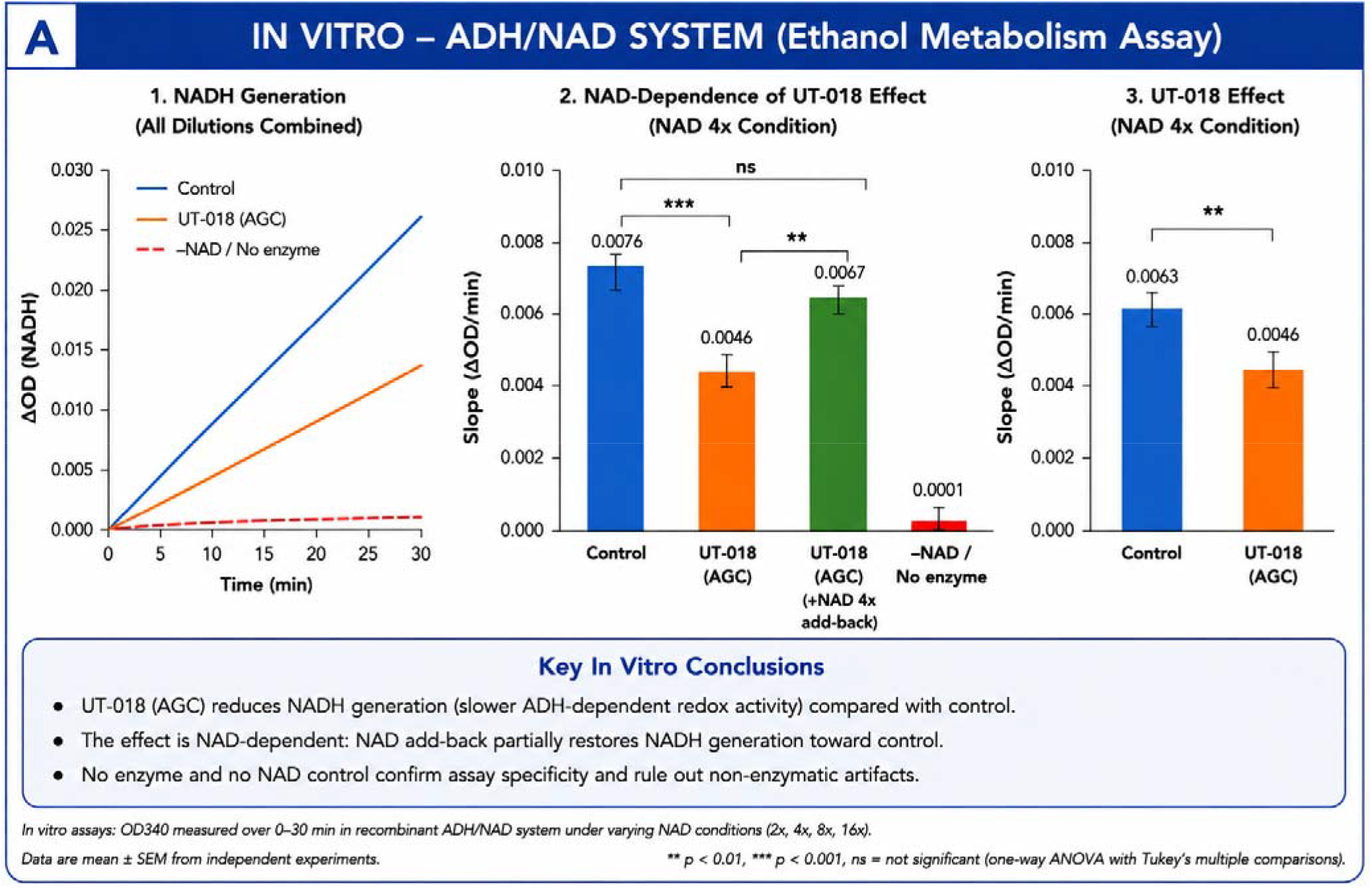
UT-018 Modulates Ethanol-Associated Redox Biology In Vitro

Effect of UT-018 on alcohol dehydrogenase-dependent NADH generation during ethanol metabolism. Recombinant ADH was incubated with ethanol, NAD^+^, and increasing concentrations of UT-018. NADH production was monitored spectrophotometrically as an indicator of ethanol-associated redox metabolism. UT-018 reduced NADH accumulation in a concentration-dependent manner, suggesting modulation of ethanol-associated redox homeostasis rather than complete inhibition of alcohol metabolism. Data are presented as mean ± SEM from independent experiments.

### UT-018 Enhances Endogenous ALDH-Associated Activity in vivo

Following acute ethanol administration in vivo, UT-018 given orally, produced a delayed increase in serum ALDH-associated activity, with the greatest response observed several hours after ethanol challenge. The response was most consistent in the mid-dose treatment(50 mM) group suggesting enhanced endogenous aldehyde detoxification capacity rather than direct enzymatic replacement. Together with the in vitro data these data also suggests that UT- 018 may modulate ethanol metabolism at multiple levels.

**Figure 2.**
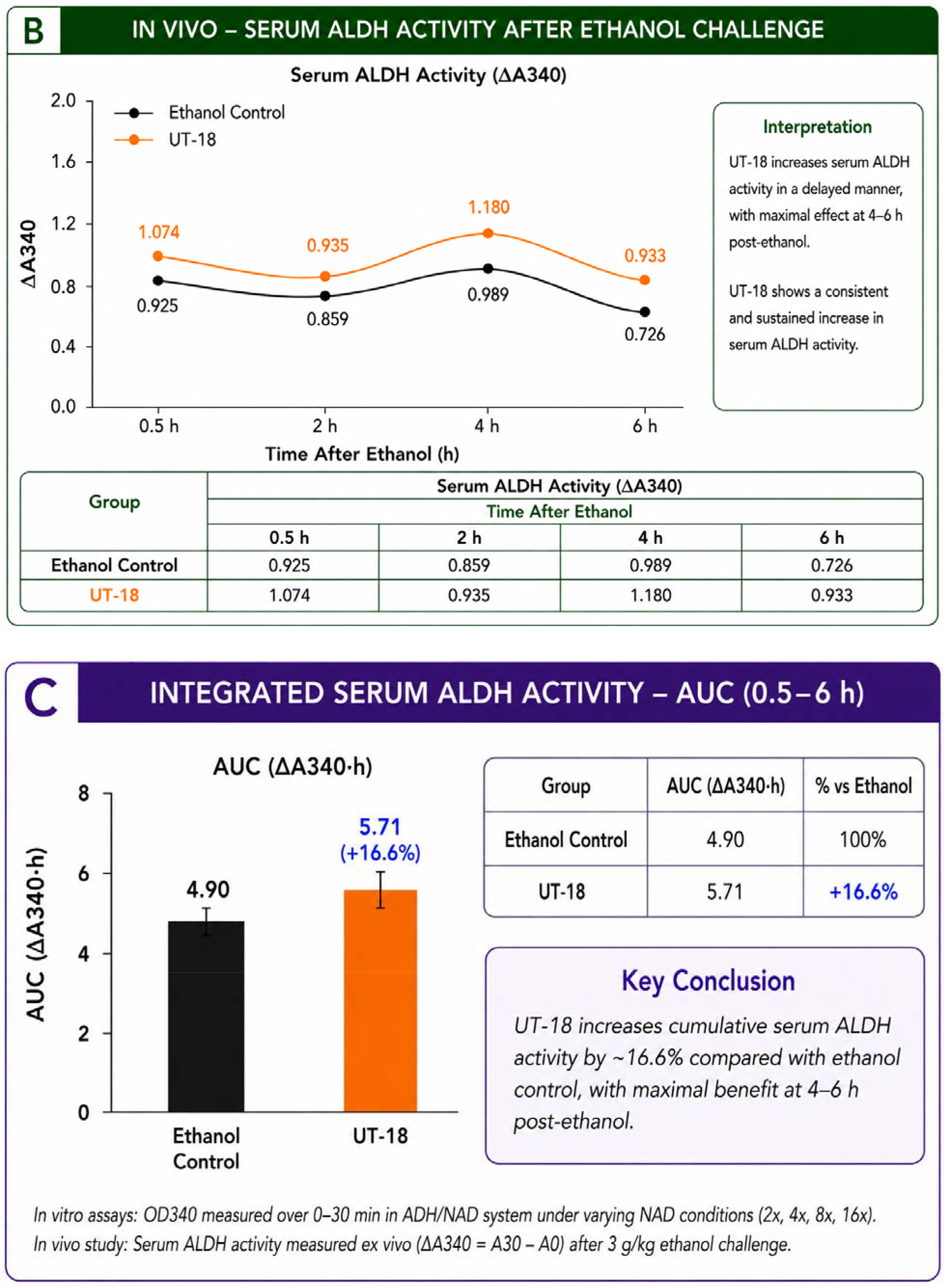
UT-018 Enhances Endogenous Aldehyde Dehydrogenase Activity Following Acute Ethanol Challenge

Time course of serum ALDH-associated activity following oral administration of UT-018 and subsequent ethanol challenge in Sprague Dawley rats. Serum samples were collected at predetermined intervals following ethanol administration and analyzed for endogenous ALDH-associated activity. UT-018 produced a delayed increase in ALDH activity, with the greatest response observed in the intermediate-dose group, consistent with enhancement of ALDH-associated detoxification. Data are presented as mean ± SEM.

### Reduction of Acetaldehyde Accumulation

Acute ethanol administration markedly increased circulating acetaldehyde levels. UT-018 oral administration treatment substantially reduced serum acetaldehyde accumulation, with the mid-dose group demonstrating the largest reduction (near normalization versus ethanol control). UT-018 produced a reduction at all doses, supporting treatment-dependent biological activity.

**Figure 3.**
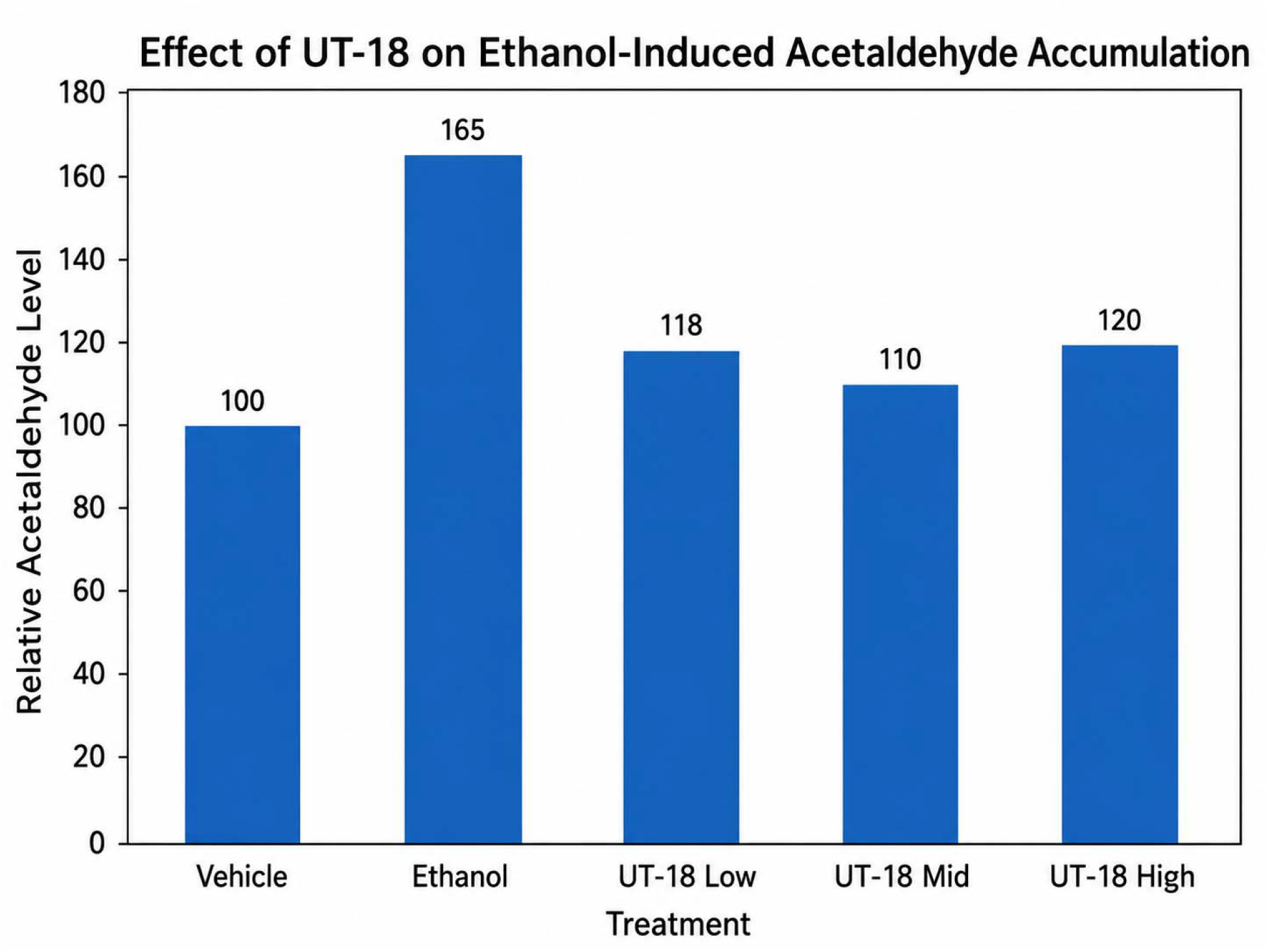
UT-018 Reduces Circulating Acetaldehyde Following Acute Ethanol Exposure

Circulating acetaldehyde concentrations following acute ethanol administration ( low 25 mM, Mid 50 mM, high 100 mM). Ethanol markedly increased systemic acetaldehyde concentrations compared with vehicle-treated animals. Treatment with UT-018 reduced acetaldehyde accumulation by approximately 27–33%. These findings support treatment-dependent enhancement of endogenous acetaldehyde detoxification. Data are expressed as mean ± SEM. Statistical significance was determined by one-way ANOVA followed by an appropriate post hoc multiple comparison test.

### Protection of Gastrointestinal Tissue

Ethanol exposure produced marked gross injury of the colon characterized by congestion and disruption of tissue architecture. UT-018 treatment reduced gross injury scores and imsupportd tissue appearance, with the low-dose formulation showing approximately 62% protection relative to ethanol controls.

Representative gross images of colonic tissue following acute ethanol administration. Ethanol-treated control animals demonstrated congestion, edema, and disruption of normal tissue architecture. Pretreatment with UT-018 imsupportd gross tissue appearance and reduced evidence of alcohol-associated injury. Gross injury scores demonstrated approximately 62% protection in the low-dose treatment group relative to ethanol controls. Images shown are representative of each treatment group.

**Figure 4.**
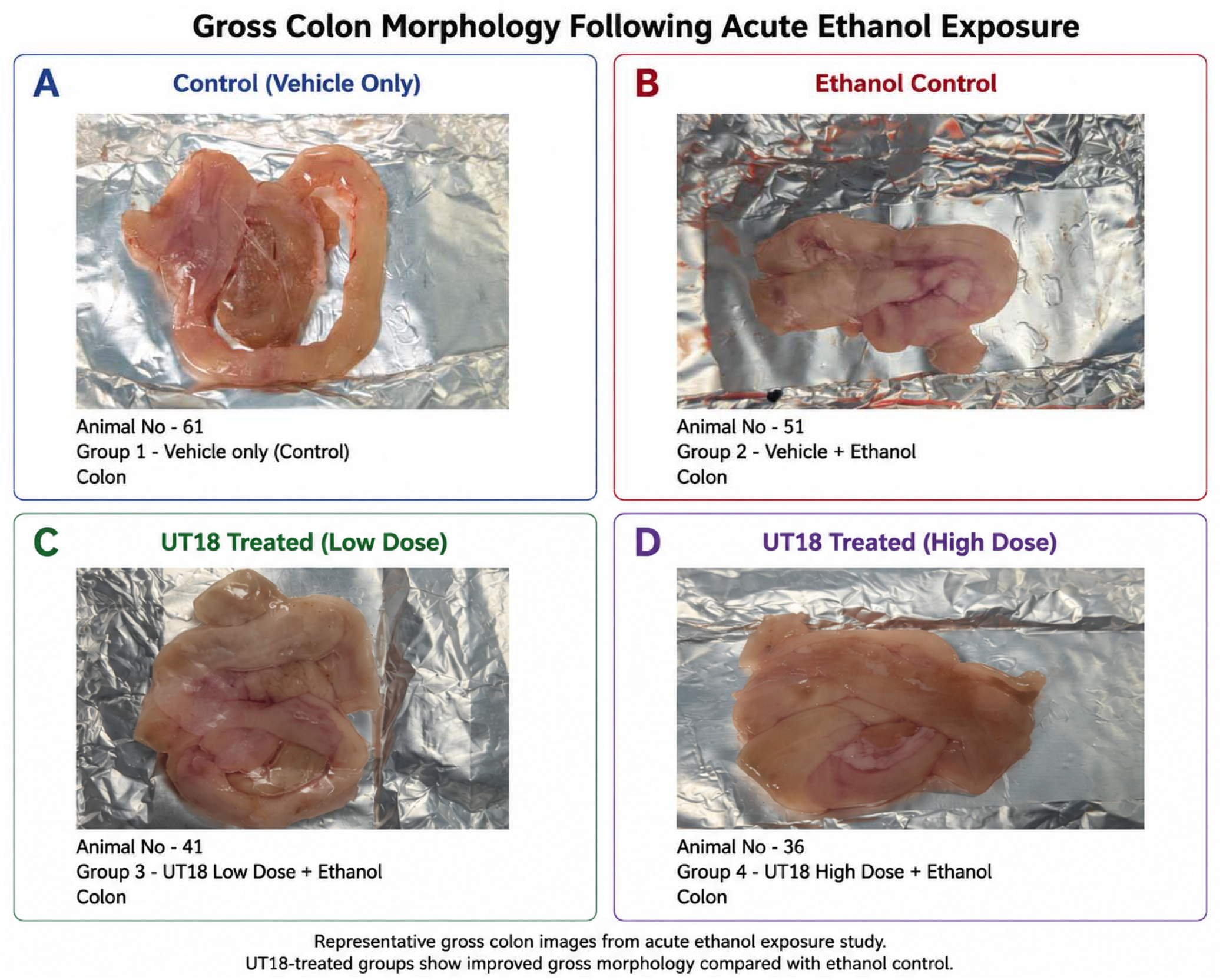
UT-018 protects against morphological changes in colon

### Preservation of Liver Morphology

Gross liver examination demonstrated modestly improved hepatic morphology in UT-018-treated animals compared with ethanol controls. The doses exhibited the more favorable liver gross morphology score, consistent with reduced hepatic injury after acute ethanol exposure.

**Figure 5.**
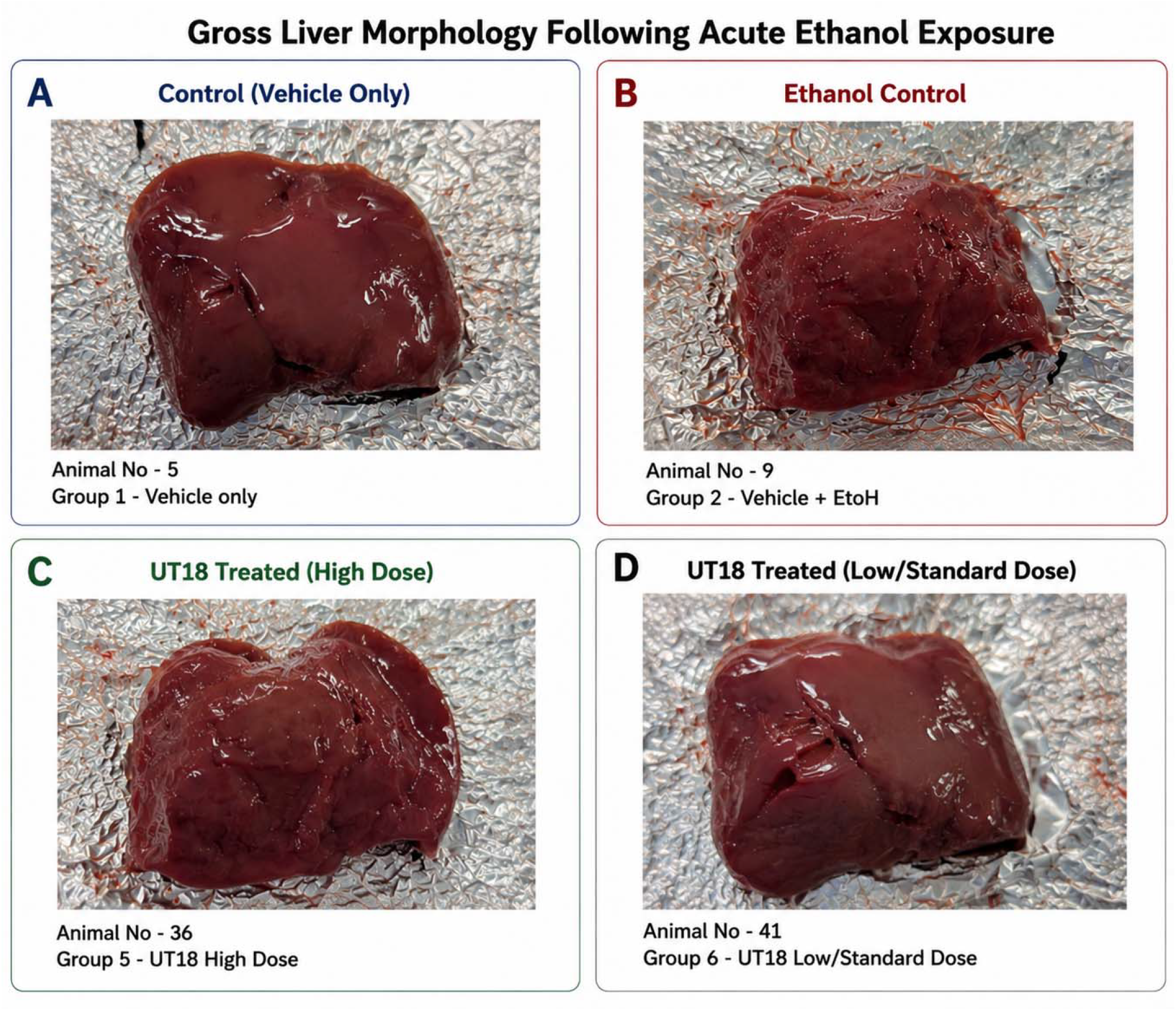
Preservation of Hepatic Morphology Following Acute Ethanol Exposure

Representative liver morphology following administration of ethanol with or without UT-018 pretreatment. Ethanol-treated control animals demonstrated macroscopic evidence of hepatic injury, including congestion and altered surface appearance. UT-018-treated animals exhibited improved hepatic morphology. Preservation of liver architecture was associated with enhancement of ALDH-associated detoxification and reduced circulating acetaldehyde concentrations.

## Discussion

Alcohol metabolism represents a tightly coordinated biochemical process in which redox balance, aldehyde metabolism, mitochondrial function, and tissue integrity are closely linked. Disruption of any one of these processes can amplify the toxic consequences of ethanol exposure.

The present studies support a model in which UT-018 acts upstream of acetaldehyde toxicity by preserving cellular redox homeostasis. Rather than directly accelerating ethanol oxidation, UT-018 appears to reduce excessive NADH accumulation, thereby maintaining a metabolic environment more favorable for endogenous ALDH function. This interpretation is supported by the NAD^+^ add-back experiments, which argue against irreversible inhibition of alcohol dehydrogenase.

An important observation is that enhanced serum ALDH-associated activity occurred several hours after ethanol exposure rather than immediately, suggesting indirect support of endogenous detoxification capacity. Such a mechanism is physiologically attractive because it augments existing metabolic pathways rather than replacing them.

The reduction in acetaldehyde accumulation translated into improvements in gastrointestinal and hepatic morphology, supporting the concept that coordinated modulation of upstream metabolic pathways can reduce downstream tissue injury. The consistent activity of UT-018 across multiple endpoints suggests that its biological effects reflect coordinated modulation of interconnected metabolic pathways.

These findings suggest that UT-018 may differ mechanistically from many currently marketed alcohol-support products, which are primarily directed toward symptomatic relief rather than correction of ethanol-associated metabolic dysfunction.

The most significant observation emerging from the present investigation is that the biological activity of UT-018 cannot be adequately explained by modulation of a single biochemical pathway. Instead, the experimental findings support a coordinated systems-level response encompassing restoration of redox homeostasis, enhancement of ALDH-associated detoxification, reduction of acetaldehyde accumulation, and preservation of gastrointestinal and hepatic integrity.

This integrated response may distinguish UT-018 from currently available interventions intended to reduce the adverse effects of alcohol consumption. Commercial products are typically designed to address dehydration, electrolyte imbalance, vitamin depletion, or oxidative stress after alcohol exposure. Although such approaches may imsupport selected symptoms, they generally do not target the fundamental metabolic disturbances responsible for alcohol-associated tissue injury.

### Redox Homeostasis Appears to Be the Primary Biological Driver

Among the various endpoints examined, modulation of cellular redox biology appears to represent the earliest measurable biological effect of UT-018.

Alcohol metabolism creates a substantial metabolic demand for NAD^+^ while simultaneously generating large quantities of NADH. This shift in the intracellular NADH/NAD^+^ ratio has profound metabolic consequences extending well beyond alcohol oxidation itself. Elevated NADH suppresses mitochondrial respiration, inhibits β- oxidation of fatty acids, reduces gluconeogenesis, alters tricarboxylic acid cycle activity, and limits oxidative phosphorylation. The resulting metabolic environment becomes progressively less favorable for efficient endogenous aldehyde metabolism.

The in vitro experiments demonstrated that UT-018 reduced excessive NADH accumulation during ethanol oxidation without abolishing alcohol dehydrogenase activity. Restoration of NADH production after supplementation with exogenous NAD^+^ provides strong evidence that the formulation acts through reversible modulation of cellular redox physiology rather than direct enzymatic inhibition. This mechanistic observation establishes a biologically plausible initiating event capable of explaining the downstream physiological responses observed throughout the study.

### Enhancement of Endogenous Aldehyde Detoxification

The delayed increase in serum ALDH-associated activity observed following administration of UT-018 provides additional evidence that the formulation enhances endogenous metabolic adaptation rather than functioning as an exogenous detoxification system.

ALDH requires adequate intracellular NAD^+^ availability to efficiently oxidize acetaldehyde to acetate. Consequently, preservation of cellular redox homeostasis would be expected to imsupport endogenous aldehyde metabolism indirectly by maintaining availability of essential metabolic cofactors.

The temporal sequence observed in the present investigation supports precisely such a mechanism. Imsupportment in redox biology preceded enhancement of ALDH-associated activity, which in turn preceded reduction of circulating acetaldehyde concentrations. This progression is entirely consistent with restoration of physiological metabolic function rather than pharmacological replacement of endogenous enzymes.

Importantly, enhancement of endogenous detoxification is likely to provide broader physiological benefits than strategies relying upon exogenous metabolic enzymes, whose activity may be limited by stability, formulation, gastrointestinal degradation, or pharmacokinetic considerations.

### Reduction of Acetaldehyde Represents the Principal Pharmacodynamic Endpoint

Acetaldehyde occupies a central position within alcohol-associated pathology because it directly mediates many of the toxic effects traditionally attributed to ethanol.

Acetaldehyde forms stable adducts with proteins, phospholipids, DNA, and cytoskeletal proteins, initiating oxidative injury, inflammatory activation, mitochondrial dysfunction, epithelial barrier disruption, and hepatocellular damage. Consequently, reduction of acetaldehyde accumulation represents one of the most biologically meaningful indicators of imsupportd alcohol metabolism.

The approximately 30% reduction in circulating acetaldehyde observed following administration of UT-018 is therefore unlikely to represent merely another biochemical measurement. Instead, it provides direct evidence that coordinated modulation of upstream metabolic pathways results in measurable reduction of exposure to one of the principal mediators of alcohol-induced toxicity.

The consistent reduction in acetaldehyde further suggests that UT-018 acts through coordinated biological activity rather than isolated modulation of a single metabolic reaction.

### Preservation of the Gut–Liver Axis

An important aspect of the present study is the simultaneous gross evaluation of both gastrointestinal and hepatic injury.

Increasing evidence indicates that alcohol-induced injury is propagated through bidirectional communication between the intestine and the liver. Ethanol disrupts epithelial barrier integrity, facilitating translocation of bacterial products and inflammatory mediators into the portal circulation. Activation of the gut–liver axis amplifies hepatic inflammation and contributes to progression of alcohol-associated liver injury.

The observation that UT-018 preserved gross morphology of both organs therefore carries mechanistic significance beyond simple tissue protection. Rather than representing independent biological events, preservation of gastrointestinal integrity and hepatic morphology likely reflects restoration of coordinated physiological function across the gut–liver metabolic axis.

Although the present investigation focused on acute ethanol exposure, preservation of this axis may have broader implications for repeated alcohol exposure and chronic metabolic injury. Future investigations incorporating histopathology, inflammatory cytokine profiling, intestinal permeability assays, and transcriptomic analyses would further define these mechanisms.

## Conclusions

The present investigation supports that UT-018 produces coordinated biological activity across multiple independent stages of alcohol metabolism. Rather than functioning as a symptomatic intervention or exogenous detoxification agent, UT-018 appears to enhance endogenous physiological resilience through restoration of cellular redox homeostasis, imsupportd aldehyde detoxification capacity, reduction of acetaldehyde accumulation, and preservation of gastrointestinal and hepatic integrity.

The consistency of the findings across biochemical, metabolic, and tissue-level endpoints supports a systems-based mechanism of action in which restoration of metabolic homeostasis serves as the initiating event leading to downstream reductions in alcohol-associated tissue injury.

Collectively, these studies provide experimental evidence that coordinated modulation of endogenous metabolic pathways represents a potential strategy for reducing the physiological consequences of acute ethanol exposure. The findings justify further preclinical investigation in chronic models of alcohol-associated injury and support clinical evaluation of UT-018 as a novel metabolic intervention for enhancing recovery following alcohol exposure.

**Figure 6.**
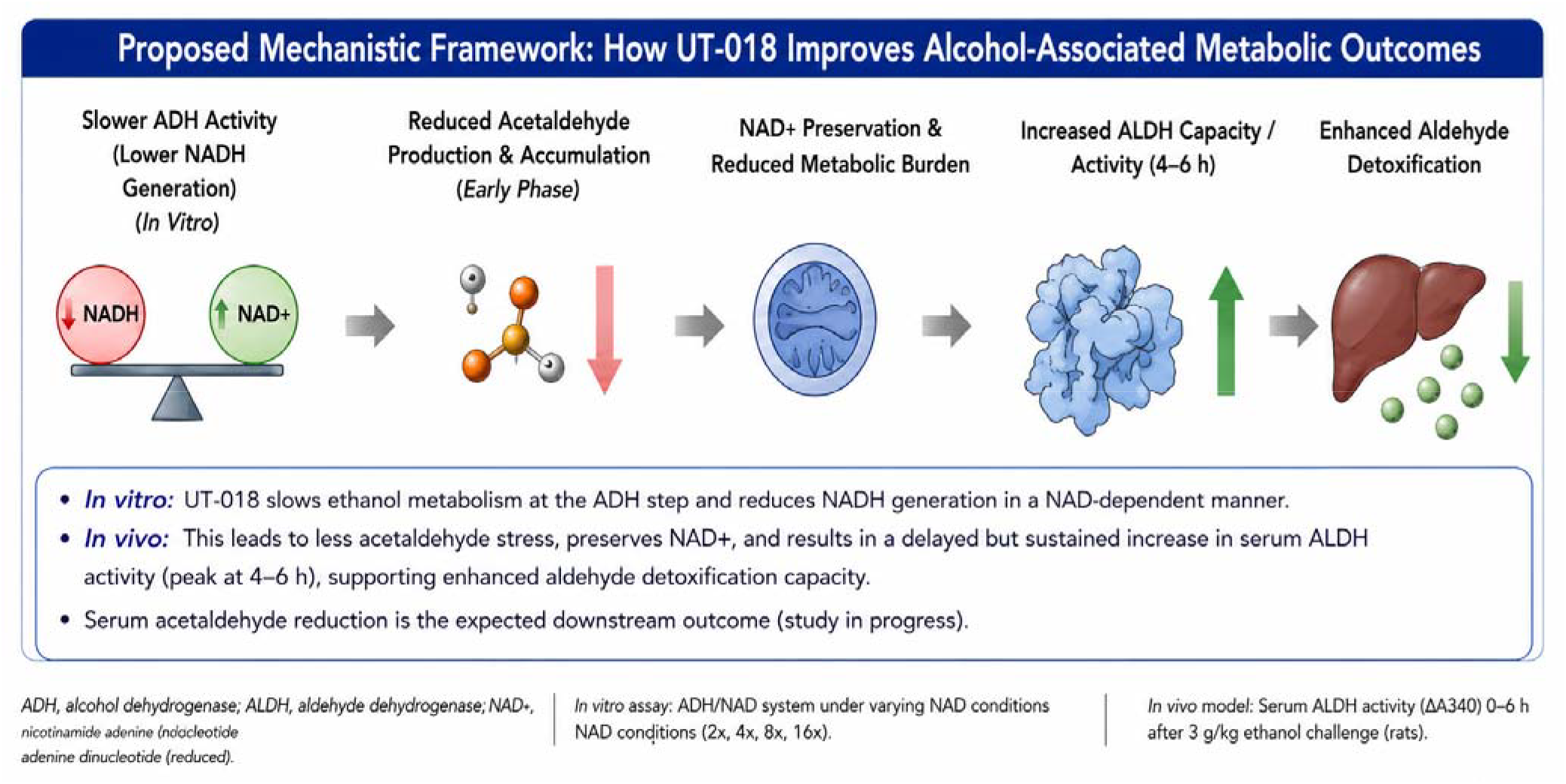
Proposed Mechanistic Framework of UT-018 Activity During Acute Ethanol Exposure

Graphical Abstract. Proposed mechanism by which UT-018 modulates alcohol-associated metabolic dysfunction. Ethanol metabolism by alcohol dehydrogenase (ADH) increases intracellular NADH and decreases NAD^+^ availability, producing redox imbalance that impairs endogenous aldehyde dehydrogenase (ALDH)-mediated acetaldehyde detoxification. Accumulation of acetaldehyde contributes to oxidative stress, gastrointestinal injury, and hepatic injury through disruption of the gut–liver axis. UT-018 is proposed to restore redox homeostasis, enhance endogenous ALDH activity, reduce circulating acetaldehyde, and preserve gastrointestinal and hepatic tissue integrity. The mechanistic model summarizes the experimental findings presented in this study and represents a working hypothesis for future investigation.

## Acknowledgements

We are grateful to Cology Biosciences, Bionest, Hyderabad, India for performing the in vivo studies.

## References

1. World Health Organization. Global Status Report on Alcohol and Health. Geneva: WHO; 2024.

2. Seitz HK, Bataller R, Cortez-Pinto H, et al. Alcoholic liver disease. Nature Reviews Disease Primers. 2018;4:16.

3. Gao B, Bataller R. Alcoholic liver disease: Pathogenesis and new therapeutic targets. Gastroenterology. 2011;141:1572–1585.

4. Cederbaum AI. Alcohol metabolism. Clinics in Liver Disease. 2012;16:667–685.

5. Zakhari S. Overview: How is alcohol metabolized by the body? Alcohol Research & Health. 2006;29:245–254.

6. Edenberg HJ. The genetics of alcohol metabolism. Alcohol Research & Health. 2007;30:5–13.

7. Crabb DW, Matsumoto M, Chang D, You M. Overview of the role of alcohol dehydrogenase and aldehyde dehydrogenase and their variants in the genesis of alcohol-related pathology. Proceedings of the Nutrition Society. 2004;63:49–63.

8. Lieber CS. Relationships between nutrition, alcohol use, and liver disease. Alcohol Research & Health. 2003;27:220–231.

9. Setshedi M, Wands JR, Monte SM. Acetaldehyde adducts in alcoholic liver disease. Oxidative Medicine and Cellular Longevity. 2010;3:178–185.

10. Brooks PJ, Enoch MA, Goldman D, et al. The alcohol flushing response: An unrecognized risk factor for esophageal cancer. PLoS Medicine. 2009;6:e1000050.

11. You M, Arteel GE. Effect of ethanol on lipid metabolism. Journal of Hepatology. 2019;70:237–248.

12. Zhong W, Zhou Z. Gut–liver axis in alcoholic liver disease. Journal of Gastroenterology and Hepatology. 2014;29(Suppl 3):11–17.

13. Albillos A, de Gottardi A, Rescigno M. The gut–liver axis in liver disease. Journal of Hepatology. 2020;72:558–577.

14. Schnabl B, Brenner DA. Interactions between the intestinal microbiome and liver diseases. Gastroenterology. 2014;146:1513–1524.

15. Bishehsari F, Magno E, Swanson G, et al. Alcohol and gut-derived inflammation. Alcohol Research. 2017;38:163–171.

16. Fernandez-Checa JC, Kaplowitz N. Hepatic mitochondrial glutathione: Transport and role in disease. Toxicology and Applied Pharmacology. 2005;204:263–273.

17. Ying W. NAD+/NADH and NADP+/NADPH in cellular functions and cell death. Antioxidants & Redox Signaling. 2008;10:179–206.

18. Verdin E. NAD+ in aging, metabolism and neurodegeneration. Science. 2015;350:1208–1213.

19. Canto C, Menzies KJ, Auwerx J. NAD+ metabolism and the control of energy homeostasis. Cell Metabolism. 2015;22:31–53.

20. Pollak N, Dölle C, Ziegler M. The power to reduce: Pyridine nucleotides—small molecules with a multitude of functions. Biochemical Journal. 2007;402:205–218.

21. He X, Sun J, Huang X, et al. Acetaldehyde-induced oxidative stress and mitochondrial dysfunction in hepatocytes. Free Radical Biology and Medicine. 2018;120:275–286.

22. Lu Y, Cederbaum AI. CYP2E1 and oxidative liver injury by alcohol. Free Radical Biology and Medicine. 2008;44:723–738.

23. Arteel GE. Oxidants and antioxidants in alcohol-induced liver disease. Gastroenterology. 2003;124:778–790.

24. Wang HJ, Gao B, Zakhari S, Nagy LE. Inflammation in alcoholic liver disease. Annual Review of Nutrition. 2012;32:343–368.

25. Szabo G, Bala S. Alcoholic liver disease and the gut–liver axis. World Journal of Gastroenterology. 2010;16:1321–1329.

26. Nagy LE. The role of innate immunity in alcoholic liver disease. Alcohol Research. 2015;37:237–250.

27. Bode C, Bode JC. Effect of alcohol consumption on the gut. Best Practice & Research Clinical Gastroenterology. 2003;17:575–592.

28. Farhadi A, Banan A, Fields J, Keshavarzian A. Intestinal barrier function and alcohol. American Journal of Gastroenterology. 2003;98:218–226.

29. Hoek JB, Pastorino JG. Ethanol, oxidative stress and cytokine-induced liver cell injury. Alcohol. 2002;27:63–68.

30. You M, Rogers CQ. Adiponectin: A key adipokine in alcoholic fatty liver. Experimental Biology and Medicine. 2009;234:850–859.

31. Lieber CS. Alcoholic fatty liver: Pathogenesis and mechanism of progression to inflammation and fibrosis. Alcohol. 2004;34:9–19.

32. Mantena SK, King AL, Andringa KK, et al. Mitochondrial dysfunction and oxidative stress in alcoholic liver disease. Mitochondrion. 2008;8:306–313.

33. Ceni E, Mello T, Galli A. Pathogenesis of alcoholic liver disease. Journal of Hepatology. 2014;61:265–280.

34. Bataller R, Brenner DA. Liver fibrosis. Journal of Clinical Investigation. 2005;115:209–218.

35. Gao B, Ahmad MF, Nagy LE, Tsukamoto H. Inflammatory pathways in alcoholic liver disease. Hepatology. 2019;69:2498–2510.

36. Jones DP. Redefining oxidative stress. Antioxidants & Redox Signaling. 2006;8:1865–1879.

37. Sies H. Oxidative stress: A concept in redox biology and medicine. Redox Biology. 2015;4:180–183.

38. Forman HJ, Zhang H. Targeting oxidative stress in disease. Nature Reviews Drug Discovery. 2021;20:689–709.

39. Chance B, Williams GR. Respiratory enzymes in oxidative phosphorylation. Advances in Enzymology. 1956;17:65–134.

40. Williamson DH, Lund P, Krebs HA. The redox state of free nicotinamide-adenine dinucleotide in the cytoplasm and mitochondria of rat liver. Biochemical Journal. 1967;103:514–527.

